# Exosome-mediated delivery of microRNAs by root-knot nematodes

**DOI:** 10.1101/2025.08.19.671007

**Authors:** M Willow H Maxwell, Alex Papp, Bharat Rohilla, Caitlin Simpson, Martin Fuller, Suruchi Roychoudhry, Chris A Bell

## Abstract

Plant-parasitic nematodes secrete molecules to manipulate their hosts, but little is known about their mode of delivery and packaging. Here, we describe microRNA-containing exosomes that are secreted by root-knot nematodes and systemically increase host susceptibility. By revealing a novel mode of nematode-plant communication, our findings outline a mechanism for the delivery of nematode patho-molecules, offering a new target for disrupting parasitism at the level of vesicle-mediated delivery.

## Introduction

Plant-parasitic nematodes pose a significant threat to agriculture in both developed and developing regions. Among them, the root-knot nematode *Meloidogyne incognita* is widely regarded as the most destructive ^1^, owing to its extremely broad host range of over 4,000 plant species ^2^. Root entry and the establishment of a complex feeding site within the host is facilitated by a suite of nematode-secreted “patho-molecules”. The secretions of juvenile nematodes are arguably the most studied and originate from two subventral gland cells (Fig. 1A), which are active pre-invasion and contribute to the degradation of the plant cell wall and suppression of host immune responses ^3–7^. Gland cell products are stored within the cytoplasm in Golgi-derived granules (700-1100 nm diameter), prior to their release from the valve ^8^. Each granule contains “minute spherical vesicles” ^8^, however their biogenesis, structure and function are unknown. An unknown neuronal cue regulates granule exocytosis and muscle contraction, which transports patho-molecules for secretion into host tissue^9.^ Animal parasitic nematodes are known to secrete exosomes that contain a range of patho-molecules, however this is not known for plant-parasitic nematode species ^10^. Over the past few decades, the plant nematology field has thoroughly detailed the repertoire and role of secreted effector proteins ^11^, however, we have little insight into the secretion of other molecules, such as nucleic acids, or whether these cargo are “naked” or packaged to aid protection/trafficking.

**Figure 1:**
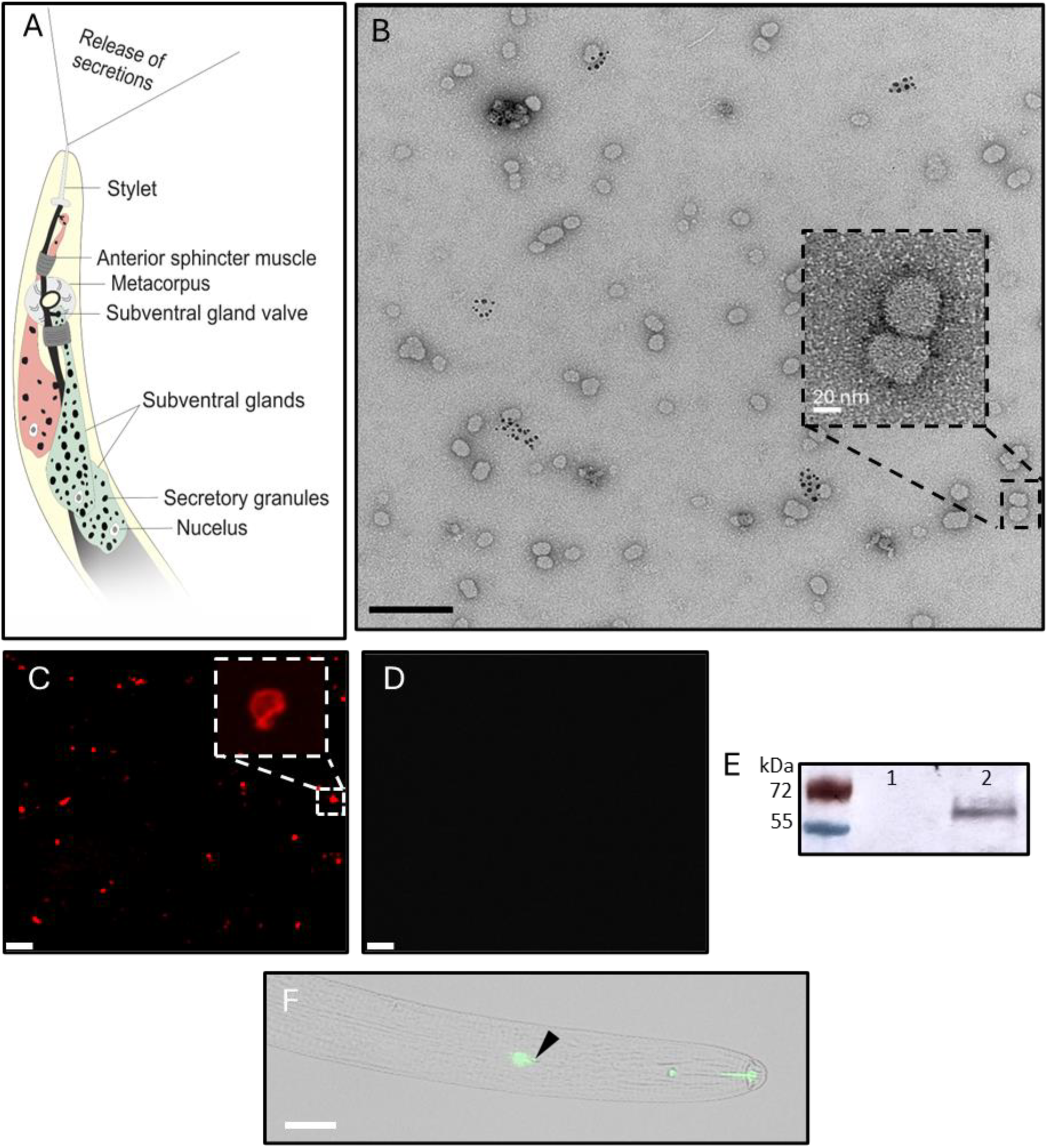
Characterisation of exosomes secreted by the plant-parasitic nematode *Meloidogyne incognita*. A) Schematic representation of the subventral gland secretion pathway of a second-stage juvenile. B) Transmission electron microscopy of resorcinol-induced secretions at 30k magnification, scale bar represents 200 nm. Inset is at 150k magnification, scale bar represents 20 nm. C) Exosomes treated with CellMask Deep Red and visualised using confocal microscopy. Scale bar represents 200 nm. D) As in C, but with 10 % Triton X-100 treatment prior to confocal microscopy. E) Western blot of the tetraspanin CD63 from non-induced (1) and resorcinol-induced secretions (2). F) Overlay images of *in situ* HCR of RAL-1 in a second-stage *M. incognita* juvenile. Metacorpus is highlighted by the black arrow.

## Results & Discussion

Subventral gland secretions are naturally induced by root exudates ^5^, however resorcinol, a neurotransmitter, triggers their release *in vitro* ^12^. Transmission electron microscopy of concentrated secretions revealed exosomes, 25-70 nm in diameter (Fig. 1B; mean 49.7 nm, SEM 0.9 nm, n = 512), which are up to 50% smaller than vesicular secretions by other nematode species ^10,13^. No exosomes were found in non-induced solutions, or the resorcinol stock.

We then investigated the structure of nematode secreted exosomes for the long-term objective of targeting their integrity, therefore disrupting the nematode’s exosome-mediated patho-molecule delivery system. CellMask staining confirmed that secreted exosomes were enclosed in lipid membranes (Fig. 1C), which commonly encapsulate cargo transported *vice versa* between pathogens and their hosts ^14^. This is supported by exosome disruption via detergent treatment (Fig. 1D). In other patho-systems, lipid-encapsulated exosomes facilitate the delivery of cytoplasmic effectors into the host cell via clathrin-mediated endocytosis ^15–17^. Exosome lipid bilayers are abundant with tetraspanins that enable membrane curvature, selection of exosome cargo and the direction and adhesion of exosomes to cell membranes ^10,18,19^. Tetraspanin CD63, a common exosome marker ^20^, was identified within secreted exosomes (Fig. 1E) (consistent with underpinning genes Minc3s00247g08503 & Minc3s01356g23074). This revealed that biological cargo from plant-parasitic nematodes is packaged in protein- and lipid-membrane bound exosomes for secretion into host roots. How the exosomes are released still remains unknown. We hypothesised that gland cell “granules” are multivesicular bodies that exocytose exosomes into the oesophagus for secretion. *M. incognita* contains orthologues of genes linked to exosome release from multivesicular bodies in *C. elegans*, such as *ral-1*, which is enriched in the subventral glands of the cyst nematode *Heterodera schachtii* (Hsc_gene_25369) ^21^. *Mi-ral-1* localises slightly to the posterior of the metacorpus of *M. incognita* second-stage juveniles. This suggests a potential role in the release of exosomes at the subventral gland valve (Fig. 1F; schematic Fig. 1A) and now requires further investigation into the release mechanism.

Secreted exosomes by other pathogens/parasites are known to contain nucleic acids ^10,22^, therefore we investigated the presence of nucleic acid-containing exosomes in *M. incognita*. Bioinformatic studies have mapped the miRNAome of a plant-parasitic nematode species, however we are still unsure whether i) miRNA are secreted, and ii) miRNA are secreted in exosomes. *M. icngontia* secreted exosomes were found to contain numerous, diverse miRNA that were enriched beyond those identified within total nematode homogenate (Fig. 2A). Highly expressed miRNA identified within induced/noninduced secretions and homogenate were removed from the list of putative secreted miRNA as possibly derived from sloughed cells (Supporting Information; e.g. developmental-regulator *let-7*). We predict that induced nematodes may have exhausted their subventral gland contents prior to nematode homogenisation, underpinning the absence of 17 secreted miRNA within the homogenate fraction.

**Figure 2:**
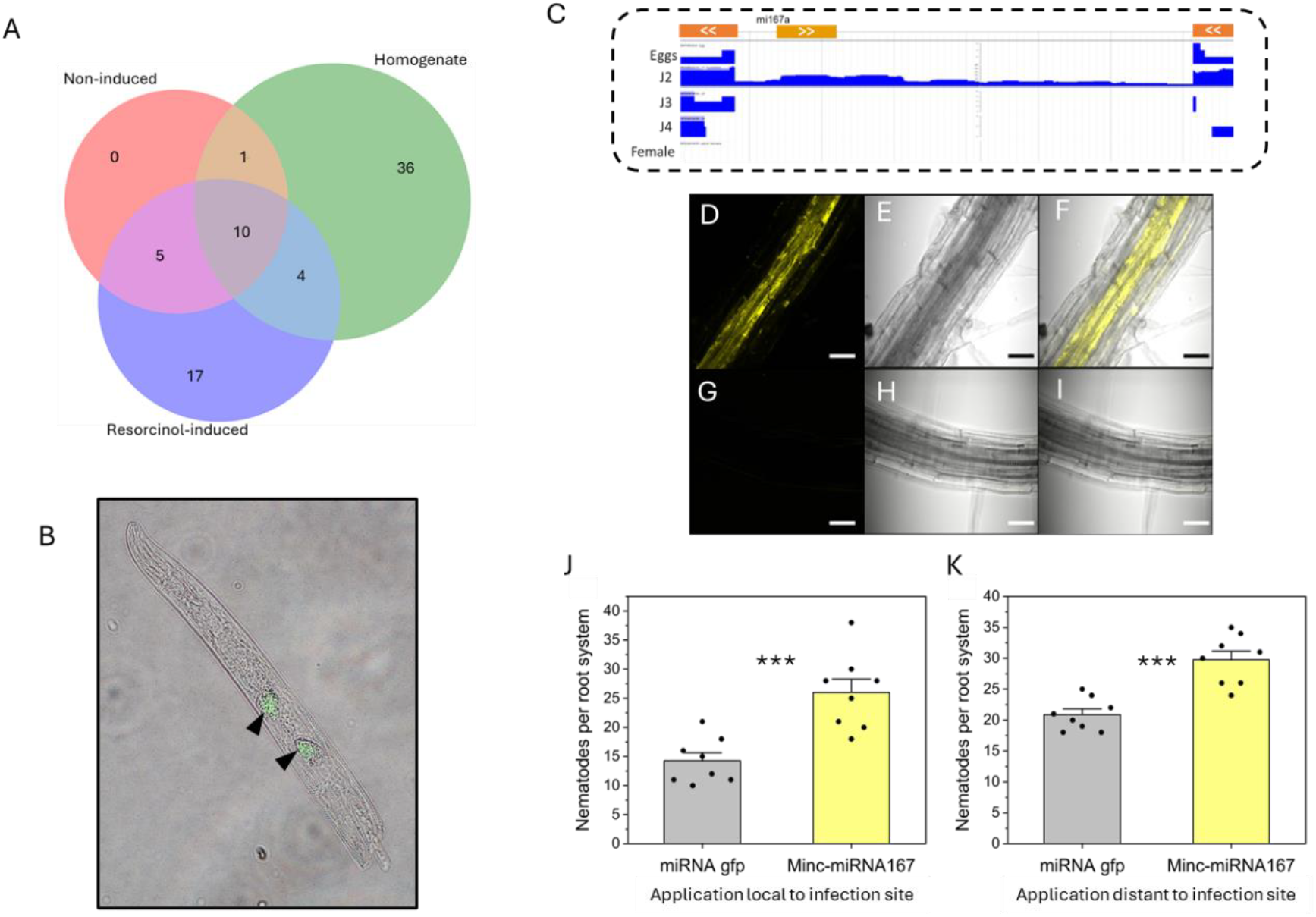
Characterisation of a miRNA present within plant-parasitic nematode secreted exosomes. A) The number of miRNA with expression counts >10 identified from sequencing of *M. incognita* (RKN) homogenate, resorcinol-induced secretions, and non-induced solutions. B) Overlay images of an *in situ* HCR of *minc-miR167a* on a second-stage *M. incognita* juvenile. Subventral glands are indicated with black arrows. C) Intronic and reverse strand location of *minc-miR167a* within the *M. incognita* genome. The outer gene exons are represented on the top row in orange, whilst *mi167a* is in yellow. Subsequent rows indicate the expression levels of each genomic region, with J2s (second-stage juveniles) uniquely expressing the intron containing *mi167a*. D, E & F) Visualisation of cy3-tagged *minc-miR167a* within the root vasculature of *Arabidopsis thaliana* Col-0, via confocal microscopy (D), with brightfield (E) and overlay (F). G, H & I) Mock treated *A. thaliana* Col-0 roots visualised by confocal microscopy. Scale bar represents 40 µm. J & K) 20 nm untagged *minc-miR167a* was applied to *M. incognita* infected root tips (J), or root tips distant to infection sites (K), every day for four days before infection was quantified. Asterisk denote significance at P<0.001 Two-sample t-test.

The most abundant miRNA enriched in nematode secretions was *miR167a*, which is widespread across plant species and regulates auxin signaling pathways that are integral to environmental outputs, including root formation and systemic resistance ^24^. *M. incognita* infection represses the plant *miR167* family at 7 & 14 dpi ^25^, however specific *miR167a* is highly abundant during early parasitism ^26^, coinciding with the activity of the subventral glands, evidencing the dynamic and context-dependent role of miRNAs seen in other systems where *miR167a/d* negatively and positively regulates immunity to fungal ^27,28^ and bacterial infection ^29^. The high expression of *miR167a* in secretions and homogenate, whilst absent within non-induced samples, led to further analyses. *In situ* hybridisation of *Mi-miR167a* confirmed its nematode origin, and expression within the subventral gland cells (Fig. 2B). Genome analysis revealed that *Mi-miR167a* is located within an intron, oriented antisense to an adjacent gene of unknown function (Fig. 2C) and has 100% sequence homology throughout the *Meloidogyne incognita* species group. This intron is only expressed in second-stage juveniles (Fig. 2C), consistent with the identification of *Mi-miR167a* within their secretions and the negative impact of host *miR167* on the development of female nematode^25^.

To determine the impact of *Mi-miR167a* within the plant, we applied synthetic Cy3-tagged *Mi-miR167a* to *Arabidopsis thaliana* roots. Systemic translocation of Cy3-tagged *Mi-miR167a* was observed via the root vasculature (Fig. 2D), in which miRNA from other pathogens is known to be transported ^30^. Application of untagged *miR167a* to infection sites increased *M. incongita* infection of tomato roots compared to infection sites treated with *gfp*-targeting miRNA (Fig. 2J). *miR167a* application at uninfected root tips distant from the nematode infection site also increased invasion of *M. incognita*, supporting a systemic effect (Fig. 2K), via vascular tissue.

In summary, we identify and characterise the exosome-based subventral gland secretion system of the root-knot nematode. We propose that secreted miRNA, and likely certain effector proteins, are released in exosomes at the gland cell valve. Exosomes likely protect the biological cargo *en route* to the host and enable their delivery into host cells via endocytosis. Here, we demonstrate that secreted-miRNAs aid systemic parasitism, potentially by targeting auxin pathways throughout the host. Our findings establish a foundation for targeting the patho-molecule delivery pathway itself, offering a broader strategy to disrupt parasitic interactions beyond single molecule approaches.

## Methods

### Induction of nematode secretions

*Meloidogyne incognita* (VW6) were maintained on tomato plants (‘Ailsa Craig’) growing in compost at 25 °C. Infective second-stage juveniles were extracted by washing the roots, cutting into 3-4 cm lengths and placing in a misting chamber. Collected juveniles were washed ten times in RNAse-free water and kept at 10 °C until use. Approximately 300 000 second-stage juvenile nematodes were placed in 1 ml 4% resorcinol for 4 hours to induce secretions. Nematodes were spun at 3000 g for 3 minutes and the supernatant was removed. The supernatant was checked to confirm no nematode contamination and either immediately used for TEM, microRNA extraction, or snap frozen at −80°C. Control nematodes were treated with RNAse free water instead of resorcinol to provide a non-induced sample.

### Transmission electron microscopy

Secretions and control samples were spun at 100 000 g for two hours ^31^ to pellet exosomes in a 30 µl fraction. Exosomes were mounted on 300 mesh, formvar/carbon coated copper grids and allowed to settle and evaporate for 20 min. Grids were washed once for 5 sec in sterile distilled water and contrasted by uranyl-acetate solution (x2 drops, 10 sec each). A FEI Tecnai G2 Spirit TEM was used to image exosome samples at a voltage of 120kV and a magnification of 30k or 150k x.

### Lipid staining

Exosomes were concentrated, as described above. Ten µl of exosomes were added to 10 µl CellMask Orange (5 µg/ml) at room temperature for 30 min. Ten µl of the labelled exosomes was analysed via a Zeiss LSM 800 confocal microscope. Upon positive CellMask Orange detection on exosomes, the reaction was duplicated to attempt to disrupt and validate the lipid membrane. After a 30 min incubation in CellMask Orange, 5 µl 10% Triton-X100 was added and incubated on ice for one hour before reimaging on the confocal microscope. Control, non-induced solutions were also observed similarly.

### Western blotting

Exosomes were mixed with 5X protein loading buffer (National diagnostics) in a total 20µl volume and denatured at 95 °C for 5 minutes. Noninduced nematode solutions were also denatured. Denatured samples were ran on a 12% gradient Mini-PROTEAN Precast gel alongside a molecular weight marker (New England biolabs Color Prestained Protein Standard, Broad Range (10-250 kDa)) at a constant voltage of 100 V for 45 minutes. Proteins were transferred onto nitrocellulose membranes using a wet transfer system (Bio-Rad) at 100 V for 75 minutes at 4 °C. Membranes were blocked in 5% (w/v) non-fat dry milk (MARVEL) and treated with Tris-buffered saline 0.1% Tween-20 (TBST) for 1 hour at room temperature to reduce non-specific binding. Membranes were incubated overnight at 4 °C with Anti-CD63 primary antibody (Abcam) diluted in 5% MARVEL-TBST with a dilution of 1:2000. Following three washes in TBST (3 × 5 minutes), membranes were incubated with Anti-rabbit secondary antibodies (1:10,000 in 5% milk/TBST) for 1 hour at room temperature. After additional TBST washes, membranes were developed overnight at 4 ° using alkaline phosphatase substrate (**SIGMAFAST™ BCIP/NBT tablets**) prepared according to the manufacturer’s instructions.

### miRNA extraction, sequencing and annotation

Secretions from approximately 300 000 resorcinol-induced *M. incognita* second-stage juveniles were collected and concentrated, as described above. Control, non-induced samples were also collected. Post-resorcinol induction, nematodes were then homogenised via micro pestle. miRNA was extracted from all samples by E.Z.N.A Micro RNA Kit (Omega Bio-tek), according to the manufacturers protocol. 50 ng of miRNA was sent to Genewiz for Small RNA-Seq, returning >20 million reads for each sample. Genewiz provided *de novo* RNA annotations as well as matches to miRbase libraries of *Caenorhabditis elegans* (model nematode species) and *Solanum lycopersicum* (tomato, host for *M. incognita*). Specifically, raw sequence reads were quality and adapter trimmed using Trimmomatic (v0.30). The trimmed reads with a length of 18 to 32 bp were retained. These trimmed reads were compared to, and annotated with, the small RNA database miRbase 22. Matched sequences were then confirmed present within the nematode genome. For novel microRNA prediction, sequences were aligned to the *M. incognita* genome and subjected to RNA folding and secondary structure analysis using miRDeep2 (v2_0_0_7). Thus, these analyses identified known and novel miRNAs. To identify putative subventral gland secreted miRNA, data were filtered to remove hits that were detected within the non-induced sample and with a read count < 10. The high expression of *miR167a* in both secretions and homogenate, yet the absence within non-induced samples, led to our further analyses on this sequence. *miR167a* was located within the *M. incognita* genome using WormBase Parasite JBrowse to determine intronic localisation and the expression patterns in each life stage using available tracks ^32^.

### In situ *hybridisation*

Nematodes were fixed and cut as previously described ^33^. Spatial gene expression profiles were determined by the hybridisation chain reaction as detailed by Sperling & Eves-van den Akker, 2023, using HCR probes produced by Molecular Instruments Inc (USA) ^34^. For miRNA *in situ* HCR a second fixation step using EDC was included to increase miRNA retention, followed by a glycine wash to quench EDC reactivity prior to probe treatments ^35^.

### Synthetic Mi-mir167a treatment

Mature *Mi-miR167a* and its complementary strand were synthesised with a 5’ CY3 tag by Eurofins Genomics to determine *in planta* transportation ^30^. Strands were annealed by combining mature miRNA and passenger strands with 5× annealing buffer (10 mM Tris, 1 mM EDTA, 50 mM NaCl in RNAse free water) for a final concentration of 2 µM. The mixture was incubated at 90 °C for 1 min, followed by a gradual decrease in temperature of 0.1°C/sec to 37 °C and held for 45 min. The roots of six-day old *Arabidopsis thaliana* were submerged in 0.2 µm CY3-Mi-miR167a for two hours, before washing three times in water. Mock water treatments were used as negative controls. Roots were visualised on a Zeiss LSM 800 confocal microscope.

Tomato ‘Ailsa Craig’ were germinated in compost before transferring to soil-free pouches. Eleven-day old seedlings were infected with approximately 100 second-stage juvenile *M. incognita*, distributed between up to four root tips. Nematodes were infected onto 1 cm square filter paper placed over root tips to avoid dispersal. Untagged *mi-miR167a* and its complementary strand were synthesised by Eurofins Genomics and annealed as described above ^35^. 20 nM *Mi-miR167a* was applied to the infected root tips each day for four days. Control plants were infected with *M. incognita* and treated with 22-nucleotide RNA complementary to *gfp*. Each treatment was replicated 12 times. At five days post infection the roots were weighed (no significant difference between treatments) and nematodes were quantified post-acid fuchsin staining. The setup was repeated but with *Mi-miR167a* applications at distant, uninfected root tips to determine systemic effects on root invasion.

## Data Availability

Micro RNA sequencing data can be found at NCBI BioProject PRJNA1271281 or in Supplementary Information.

## Acknowledgments

C.A.B. is supported by a BBSRC Discovery Fellowship (BB/X009823/1) and Michael Beverley Innovation Fellowship. We thank Prof P Urwin for hosting these fellowships in his laboratory. We thank University of Leeds for supporting M.W.H.M, BBSRC IAA for supporting B.R (127410) and BSPP for supporting A.P. For the purpose of open access, the author has applied a Creative Commons Attribution (CC BY) licence to any Author Accepted Manuscript version arising from this submission.

